# Laminar-specific cortico-cortical loops in mouse visual cortex

**DOI:** 10.1101/773085

**Authors:** Hedi Young, Beatriz Belbut, Margarida Baeta, Leopoldo Petreanu

## Abstract

Many theories propose recurrent interactions across the cortical hierarchy, but it is unclear if cortical circuits are selectively wired to implement looped computations. Using subcellular channelrhodopsin-2-assisted circuit mapping in mouse visual cortex, we compared feedforward (FF) or feedback (FB) cortico-cortical input to cells projecting back to the input source (looped neurons) with cells projecting to a different cortical or subcortical area (non-looped neurons). Despite having different laminar innervation patterns, FF and FB afferents showed similar cell-type selectivity, making stronger connections with looped neurons versus non-looped neurons in layer (L) 5 and L6, but not in L2/3. FB inputs preferentially innervated the apical tufts of looped L5 neurons, but not their perisomatic dendrites. Our results reveal that interareal cortical connections are selectively wired into monosynaptic excitatory loops involving L6 and the apical dendrites of L5 neurons, supporting a role of these circuit elements in hierarchical recurrent computations.

## Introduction

The complex network of cortical areas can be hierarchically ordered based on the anatomy of interareal cortico-cortical (CC) projections. Lower areas send bottom-up “feedforward” (FF) inputs to higher areas, which reciprocate with anatomically distinct top-down “feedback” (FB) inputs ^1,2^. The fact that a similar hierarchical architecture is observed across areas and across species, from rodents to humans, suggests that FF and FB interactions reside at the core of cortical function. As with local cortical circuits, posited to implement a conserved canonical computation in different cortical areas ^3,4^, long-range cortical connections could be performing stereotyped functions in different areas and in different species. However, the precise function of FF and FB interactions in hierarchical processing remains poorly understood.

Several theories of hierarchical computation involve looped interactions between areas, in which FF and FB pathways selectively influence each other in a bidirectional manner ^5–11^. While it is well established that cortical areas are densely interconnected and that FF inputs are always reciprocated by FB inputs ^1,12–14^, selective engagement of CC inputs with neurons projecting back to the source of those inputs (looped neurons) would require precise wiring specificity, since these neurons are closely intermingled with neurons projecting elsewhere (non-looped neurons). One possibility is that CC projections specifically modulate looped neurons indirectly via intermediary inhibitory or excitatory cells in the local circuit. Another possibility, not mutually exclusive with the previous one, is that CC projections selectively synapse onto looped neurons directly to form interareal monosynaptic loops, which would be excitatory since most long-range cortical afferents are glutamatergic. However, whether CC connections are wired to selectively facilitate looped computations remains unknown. Cortical projection neurons can be divided into 3 broad classes: intratelencephalic (IT) neurons, which project to cortical areas, pyramidal tract (PT) neurons, which project to multiple subcerebral areas including the midbrain, and corticothalamic (CT) neurons, which project predominantly to the thalamus ^4,15^. Thus, long-range CC projections could selectively participate in excitatory monosynaptic loops by preferentially contacting looped IT neurons, while avoiding neighboring non-looped IT, PT and CT neurons.

Here we measured the connectivity of CC afferents to different types of projection neurons in mouse visual cortex to test whether they are wired to specifically engage in monosynaptic looped interactions. Using a combination of subcellular channelrhodopsin-2(ChR2)-assisted circuit mapping (sCRACM) ^16^ and injections of multiple retrograde tracers, we found that FF and FB axons preferentially innervate looped neurons in layer (L) 5 and L6, but not in L2/3. Thus, both ascending and descending hierarchical streams display the same selectivity for specific looped projection neurons despite their different anatomical profiles. Moreover, preferential innervation of looped L5 neurons often involved synapses made on their apical, but not basal, dendrites. Our results suggest a prevalent role of excitatory looped interactions in hierarchical computation and identify layers and dendritic compartments that might be mediating them.

## Results

### Neurons with different projection patterns are intermingled in visual areas

Primary visual cortex (V1), the lowest-order area of mouse visual cortex, sends FF projections to the higher-order lateromedial (LM) and anteromedial (AM) visual areas, which in turn reciprocate with FB connections ^17–19^. Using dual injections of retrograde tracers, we measured the laminar distribution of different projection neurons in V1 and in LM (Figure 1). In each experiment, we compared the laminar distribution of LM- or V1-projecting IT neurons with either IT neurons projecting to AM, PT neurons projecting to the superior colliculus (SC), or CT neurons projecting to the dorsal lateral geniculate nucleus (dLGN) or to the lateral posterior nucleus (LP) of the thalamus. In each case, the different projection neurons were closely intermingled (Figure 1A,B). In both V1 and LM, IT neurons were distributed across all layers except L1, including L4 ^19,20^, indicating that FF and FB projections originate from neurons spanning most of the cortical depth. In contrast, PT and CT neurons were confined to L5 and L5/6, respectively, as previously described ^4^ (Figure 1A,B). In both V1 and LM, we found double-labeled IT neurons in L2–6 after injecting tracers in two different cortical areas, indicating that subpopulations of ascending and descending projection neurons have diverging axons innervating more than one visual cortical area ^21^. However, IT neurons were rarely double-labeled when tracers were injected in a cortical and subcortical area, confirming that cortical- and subcortical-projecting neurons constitute different classes of projection neurons ^4,22,23^. Using adeno-associated virus encoding green fluorescent protein (AAV-GFP), we anterogradely traced V1→LM FF axons and LM→V1 FB axons and measured their laminated termination pattern (Figure 1C,D). FF axons in LM were present in all layers but were denser in L2/3 and L6. FB axons in V1 arborized in L1 and L6, while avoiding middle layers. Given the laminar distribution of the different projection neurons and the termination pattern of CC afferent axons, FF and FB projections could potentially directly innervate both looped and non-looped projection neurons in each cortical layer (Figure 2A). Moreover, the proximity of looped and non-looped cells indicates that FF and FB axons are equally accessible to them. Thus, functional mapping of FF and FB connections is required to reveal any underlying synaptic specificity, since axo-dendritic overlap does not always predict connectivity ^4^.

**Figure 1.**
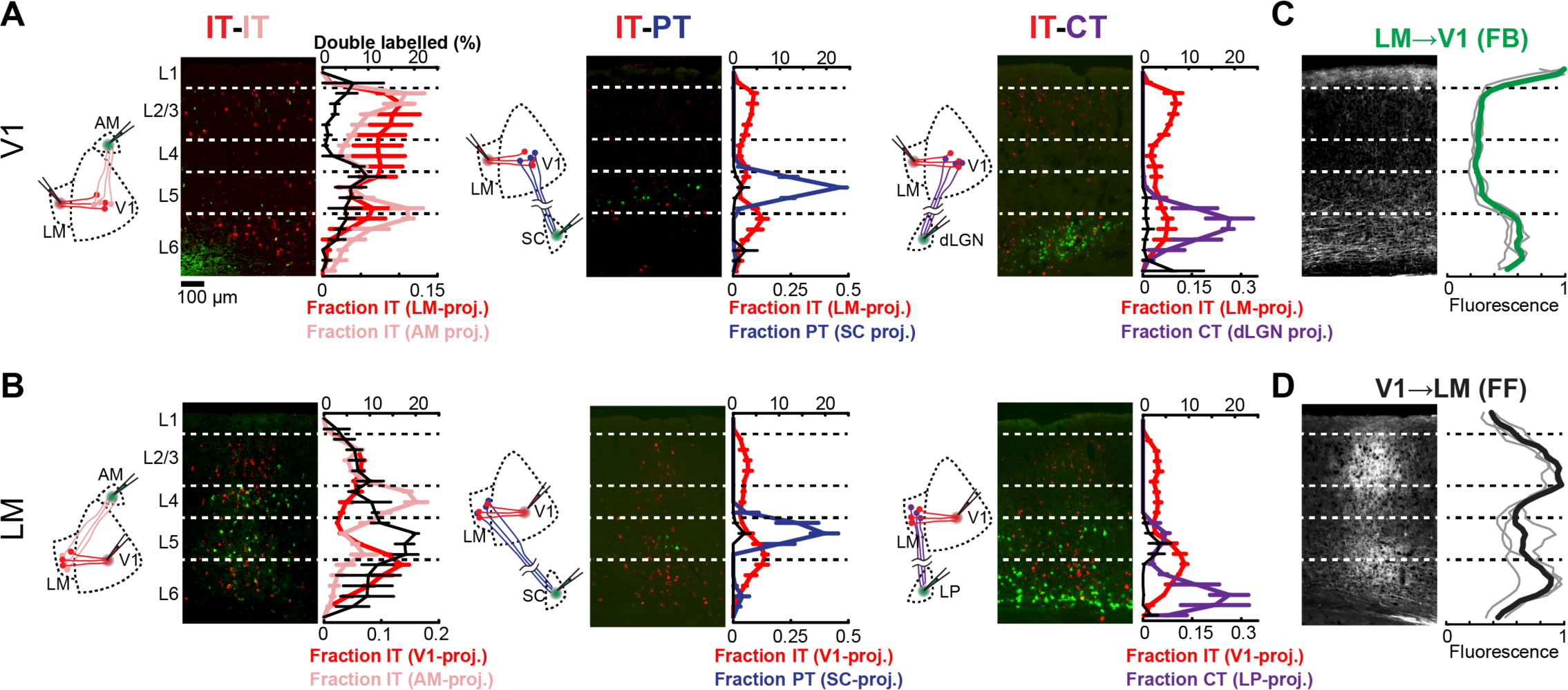
Cortical neurons projecting to different areas are intermingled and accessible to FF and FB axons. **A**, Distribution of retrogradely labeled projection neurons in V1 after injection of a red-fluorescent tracer in LM and an infrared-fluorescent tracer in either AM, SC or dLGN. Left, experimental configuration; Center, representative fluorescent histological section, with infrared fluorescence shown in green; Right, laminar distribution of the different projection neurons (n=3 animals per group; black trace, double-labeled neurons). **B**, Distribution of retrogradely labeled projection neurons in LM after injection of a red-fluorescent tracer in V1 and an infrared-fluorescent tracer in either AM, SC or LP (n=3 animals per group). **C**, Distribution of anterogradely labeled LM FB axons in V1. Left, representative fluorescent histological section; Right, axonal fluorescence across cortical depth. Individual mice, thin gray traces; average, thick green trace (n=3 animals). **D**, Distribution of anterogradely labeled V1 FF axons in LM.

**Figure 2.**
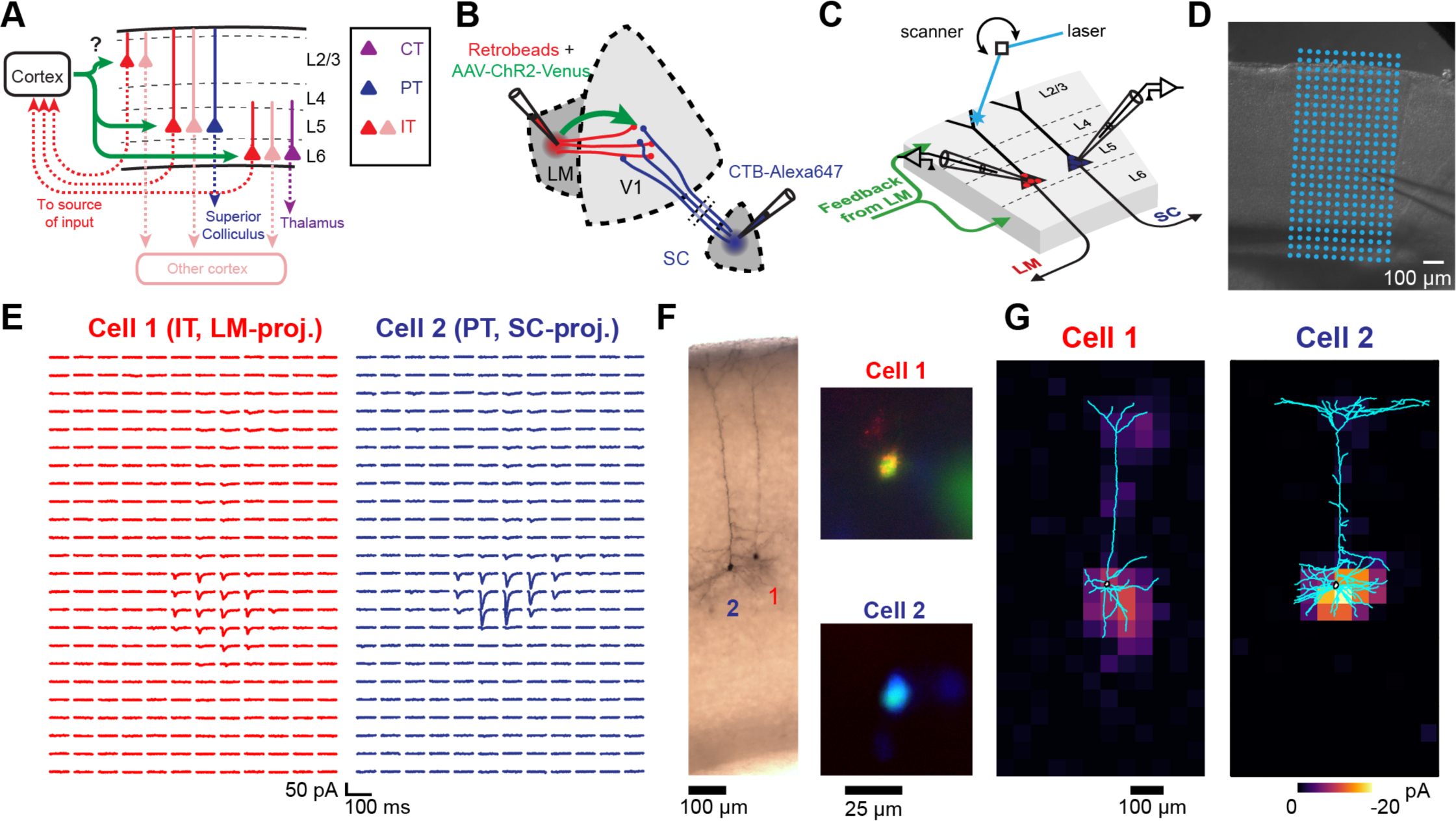
Measuring the strength and dendritic distribution of CC inputs to different projection neurons. **A**, We probed the strength of CC inputs to looped and non-looped neurons in different cortical layers. **B**, Example experiment configuration. Retrograde tracers are injected in two areas to label different projection neurons. One cortical area is also co-injected with AAV-ChR2-Venus to express ChR2 in a specific CC projection. **C**, Example sCRACM experiment. Pairs of neighboring retrogradely labeled neurons in the same cortical layer were sequentially recorded. During each recording, a laser beam was scanned over the dendrites of the cell at different locations in a grid pattern. **D**, Brightfield image of an acute coronal cortical slice showing the recording pipette and photostimulation grid. **E**, Excitatory postsynaptic currents (EPSCs) recorded from a pair of neighboring L5 neurons, evoked by photostimulating ChR2+ LM→V1 FB axons on a grid. **F**, Left, dendritic morphology staining of the recorded pair. Right, identity of the recorded projection neuron was confirmed by fluorescence in the soma of both a retrograde tracer and a different-colored dye introduced from the internal patch pipette solution. **G**, sCRACM maps of the recorded pair overlaid on their reconstructed dendrites. Non-zero EPSCs are color-coded to represent mean amplitude.

### Mapping the cell-type specificity of CC connections

To measure the cell-type selectivity of CC connections, we combined sCRACM ^16^ with multiple injections of fluorescent retrograde tracers (Figure 2B). We injected AAV-ChR2 mixed with a retrograde tracer in either V1, LM or AM to express ChR2 in FF or FB axons and to retrogradely label looped IT neurons projecting back to the source of ChR2-expressing (ChR2^+^) axons. We also injected a different retrograde tracer in either a second cortical area, the superior colliculus or the thalamus to label non-looped IT, PT or CT neurons (Figure 2A). We recorded from pairs of neighboring neurons in the same cortical layer in either V1 or higher-order visual areas LM or AM in acute brain slices containing FB or FF ChR2^+^ axons, respectively. For each pair, one cell projected to the source of ChR2^+^ inputs and one cell projected to a different cortical or subcortical area, as indicated by the retrograde tracer type in the soma (Figure 2C,F). During the recording, we used galvanometer mirrors to rapidly photostimulate ChR2^+^ axons with a blue laser at different locations around the cell to evoke excitatory postsynaptic currents (EPSCs) in the presence of sodium channel blocker tetrodotoxin (TTX; Figure 2C–E). This provided a measure of both the strength and location of monosynaptic CC inputs. We then compared the strength of monosynaptic FF or FB inputs in defined dendritic compartments in pairs of neurons projecting to different areas (Figure 2E–G)^16,17,24,25^.

### FF and FB inputs innervate specific dendritic compartments of projection neurons

We detected monosynaptic FF and FB inputs in every cell type and analyzed their dendritic distribution (Figure 3). Individual sCRACM maps were normalized to their maximum response, aligned relative to pia and averaged. They thus represent the relative distribution of CC inputs within the dendritic tree for each class of projection neuron. In L2/3, FF inputs were largely confined to perisomatic dendrites. In L6, while FF connections also targeted perisomatic dendrites in both CT and IT neurons, we could detect additional FF input on the apical dendrites of IT neurons extending across L5 (Figure 3A, Figure 4B). Similarly, FF input contacted L5 IT neurons in both the perisomatic dendrites and along their apical dendritic trunk spanning L4 and L2/3 (Figure 3A, Figure 5B). Thus, like other long-range inputs ^16^, FF axons innervate several dendritic domains of L5 pyramidal neurons, revealing that targeting of apical dendrites is not an exclusive property of FB axons (see below).

**Figure 3.**
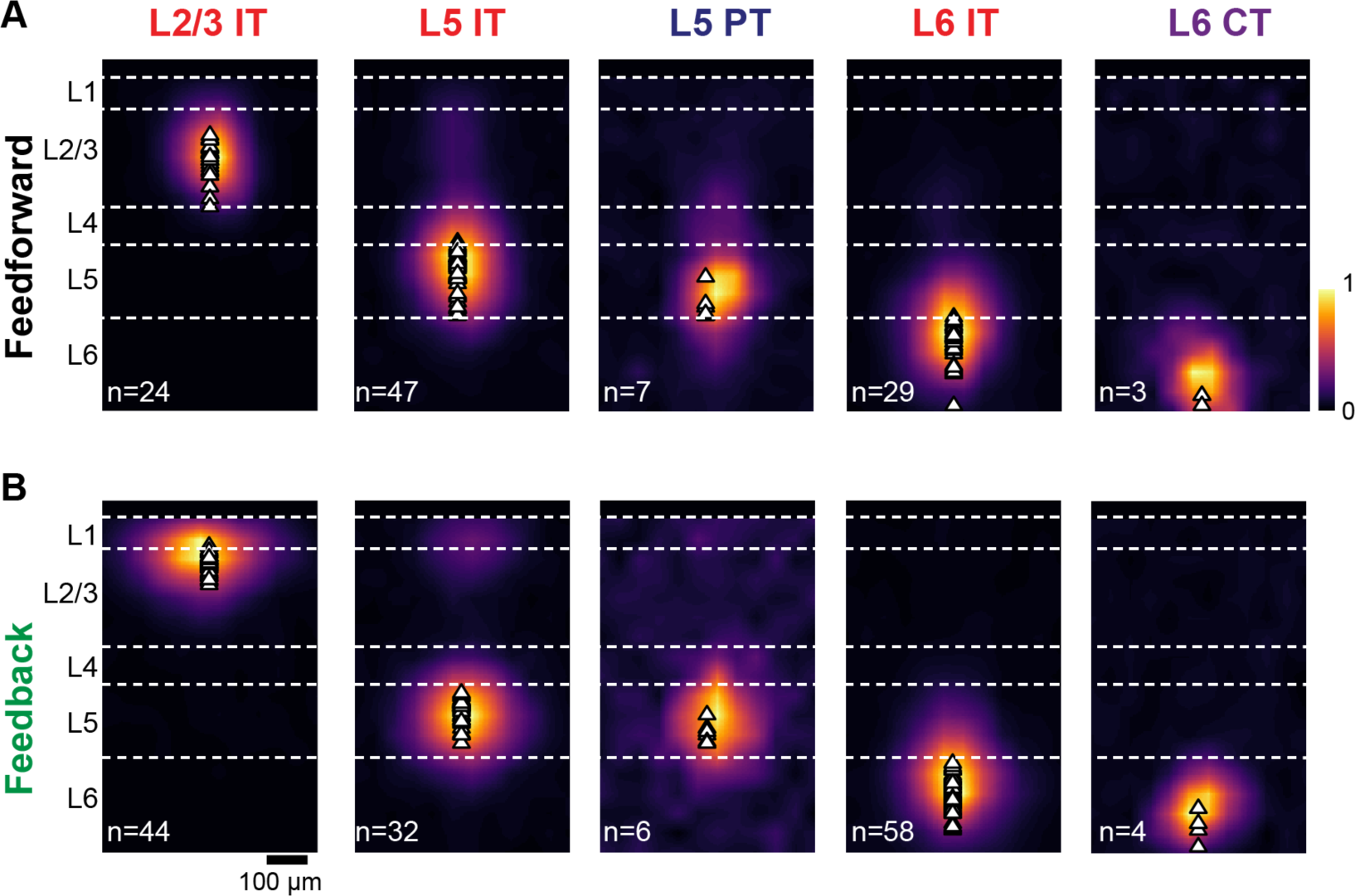
Dendritic distribution of FF and FB inputs to different projection neuron classes. **A**, Group averages of sCRACM maps aligned by pia position showing FF input to the different cell types. Triangles, soma position. **B**, Group averages of sCRACM maps showing FB input to the different cell types.

**Figure 4.**
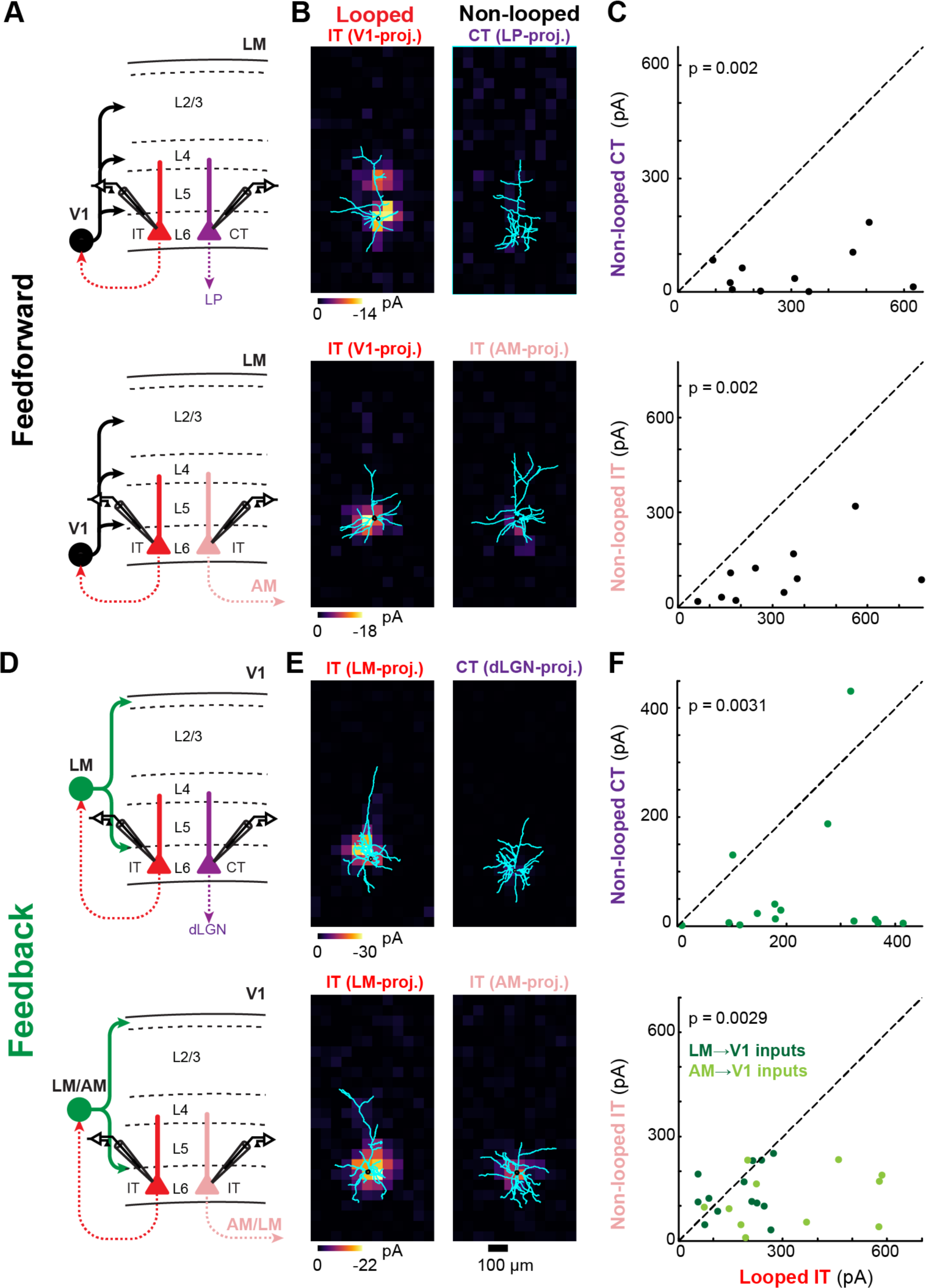
Both FF and FB inputs preferentially innervate looped IT neurons over neighboring non-looped CT or IT neurons in L6. **A**, Configuration of experiments comparing strength of V1 FF input to pairs of different L6 projection neurons in LM. Top, looped IT neuron vs. non-looped CT neuron. Bottom, looped vs. non-looped IT neuron. **B**, Example pairs of sCRACM maps overlaid on reconstructed dendrites. Each pair shows monosynaptic V1 FF inputs to a looped neuron (left) and an adjacent non-looped neuron (right) recorded in L6 of LM. **C**, Top, paired comparisons of total FF input to looped IT neurons vs. non-looped CT neurons in L6 (n=10 cell pairs); Bottom, paired comparisons of total FF input to looped vs. non-looped IT neurons in L6 (n=10 cell pairs). Statistics, two-tailed Wilcoxon signed-rank test for paired samples. **D**, Configuration of experiments comparing strength of LM or AM FB input to pairs of different L6 projection neurons in V1. Top, looped IT neuron vs. non-looped CT neuron. Bottom, looped vs. non-looped IT neuron. **E**, Example pairs of sCRACM maps overlaid on reconstructed dendrites. Each pair shows monosynaptic LM FB inputs to a looped neuron (left) and an adjacent non-looped neuron (right) recorded in L6 of V1. **F**, Top, paired comparisons of total FB input to looped IT neurons vs. non-looped CT neurons in L6 (n=14 cell pairs); Bottom, paired comparisons of total FB input to looped vs. non-looped IT neurons in L6 (n=25 cell pairs; dot colors denote the photostimulated inputs: LM→V1, n=14; AM→V1, n=11).

**Figure 5.**
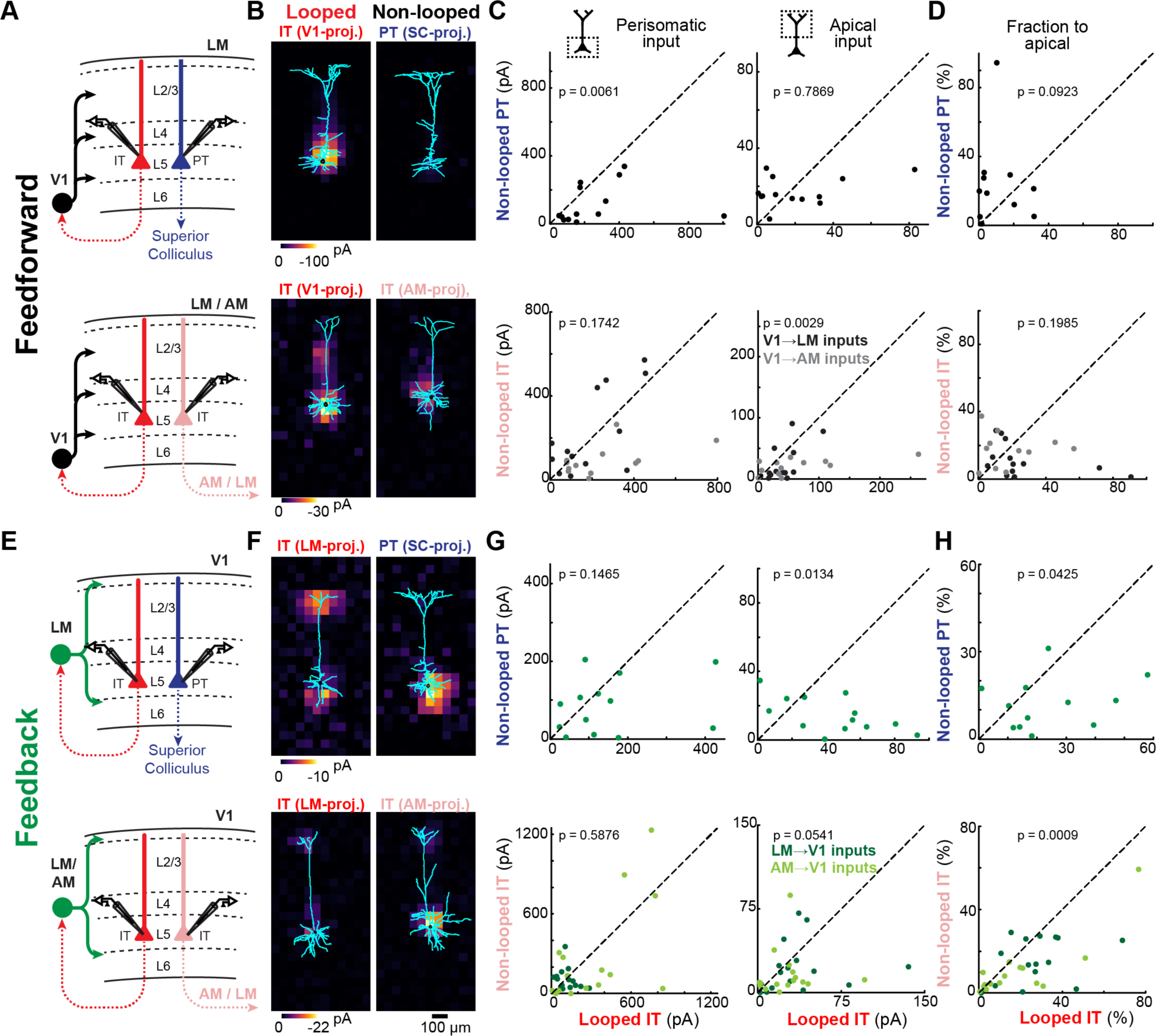
Both FF and FB connections selectively target looped L5 neurons. **A**, Configuration of experiments comparing strength of V1 FF input to pairs of different L5 projection neurons in LM or AM. Top, looped IT neuron vs. non-looped PT neuron. Bottom, looped vs. non-looped IT neuron. **B**, Example pairs of sCRACM maps overlaid on reconstructed dendrites. **C**, Top left, paired comparisons of perisomatic FF input to looped IT neurons vs. non-looped PT neurons in L5 (n=13 pairs); Bottom left, paired comparisons of perisomatic FF input to looped vs. non-looped IT neurons in L5 (n=25 pairs; dot colors denote the photostimulated inputs: V1→LM, n=13; V1→AM, n=12). Right, paired comparisons of apical dendritic FF input in the same cell pairs. Statistics, two-tailed Wilcoxon signed-rank test for paired samples. **D**, Paired comparisons of the fraction of total FF input impinging on apical dendrites. Only pairs in which both cells showed detectable input (total input > 5 pA) were included (top, n=12 pairs; bottom, n=24 pairs, V1→LM inputs, n=13, V1→AM inputs, n=11). **E**, Configuration of experiments comparing strength of LM or AM FB input to pairs of different L5 projection neurons in V1. Top, looped IT neuron vs. non-looped PT neuron. Bottom, looped vs. non-looped IT neuron. **F**, Example pairs of sCRACM maps overlaid on reconstructed dendrites. **G**, Top left, paired comparisons of perisomatic FB input to looped IT neurons vs. non-looped PT neurons in L5 (n=13 pairs); Bottom left, paired comparisons of perisomatic FB input to looped vs. non-looped IT neurons in L5 (n=32 pairs; dot colors denote the photostimulated inputs: LM→V1, n=16; AM→V1, n=16). Right, paired comparisons of apical dendritic FB input in the same cell pairs. **H**, Paired comparisons of the fraction of total FB input impinging on apical dendrites (top, n=13 pairs; bottom, n=31 pairs, LM→V1 inputs, n=16, AM→V1 inputs, n=15).

As with FF inputs, FB afferents innervated the perisomatic compartments of all recorded cell types. However, unlike FF inputs, a significant fraction of FB input to L2/3 and L5 neurons was located on their distal dendrites in L1 (Figure 3B), consistent with previous measurements ^16,24^ and with the laminar profile of FB axons (Figure 1C). Apical tuft inputs were more readily detected in L5 IT neurons than in PT neurons, suggesting differential innervation by FB fibers (Figure 2G, Figure 3B and Figure 5F, see below).

### CC inputs selectively innervate looped L6 neurons

We first measured the connectivity of FF and FB inputs to L6 neurons projecting to different areas. We compared the input strength to looped IT neurons (i.e. neurons projecting back to the FF or FB input source) versus either neighboring non-looped IT neurons projecting to another cortical area or non-looped CT neurons (Figure 4). When measuring V1 FF inputs to L6 neurons in LM (Figure 4A–C), total input to looped IT neurons was consistently stronger than to non-looped IT or CT neurons (median ratio of input; looped IT neuron/CT neuron: 7.2, p=0.002; looped IT neuron/non-looped IT neuron: 3.6, p=0.002). Likewise, FB inputs to V1 also selectively targeted L6 looped neurons (Figure 4D–F). LM-projecting neurons received ~10-fold stronger inputs from LM compared to neighboring dLGN-projecting CT neurons (median ratio of input; looped IT neuron/CT neuron: 9.6, p=0.0031). For the looped versus non-looped comparison among IT neurons, we measured FB input in V1 from two different sources, LM and AM. In both cases, FB preferentially connected to IT neurons projecting back to the source of FB, but only AM FB reached statistical significance (Figure 4F; median ratio of input; AM FB, looped IT neuron/non-looped IT neuron: 3.08, p=0.0049; LM FB, looped IT neuron/non-looped IT neuron: 1.21, p=0.1530). Considering both sources of FB together, looped IT neurons received ~1.6-fold more input than adjacent non-looped IT neurons (median ratio of input; AM & LM FB combined, looped IT neuron/non-looped IT neuron: 1.6, p=0.0029). Thus, both FF and FB axons form stronger connections with looped L6 neurons when compared to non-looped L6 neurons projecting either to a different cortical area or to a thalamic nucleus.

### FB inputs selectively innervate the apical dendrites of looped L5 neurons

We next asked whether CC inputs form distinct connectivity patterns with different L5 pyramidal neurons (Figure 5). As FF and FB inputs innervated L5 neurons in both perisomatic and apical dendritic regions (Figure 3, Figure 5B,F), we analyzed input strength in these two innervation domains separately. First, we compared V1→LM FF inputs to looped IT neurons projecting back to V1 versus non-looped PT neurons projecting to SC (Figure 5A). When considering input to all dendritic compartments, FF fibers preferentially innervated looped IT neurons over PT neurons (Figure 5B; median ratio of input; looped IT neuron/PT neuron: 1.66, p=0.0105). This difference was due to stronger perisomatic inputs to looped neurons, since no difference was observed in apical inputs (Figure 5C,D). Second, when compared to neighboring non-looped IT neurons, looped neurons in AM received stronger FF inputs on both their perisomatic and apical dendrites (median ratio of input; V1→AM, looped IT neuron/non-looped IT neuron: perisomatic, 2.05, p=0.0024; apical, 2.4, p=0.0268), but looped neurons in LM showed a tendency to receive stronger FF inputs on their apical dendrites only (median ratio of input; V1→LM, looped IT neuron/non-looped IT neuron: perisomatic, 0.89, p=0.4973; apical, 2.36, p=0.0681). Thus, when taking V1→AM and V1→LM together, FF inputs to L5 IT neurons showed a robust preference for looped neurons in their apical compartments (Figure 5B–D; median ratio of input; looped IT neuron/non-looped IT neuron: perisomatic, 1.43, p=0.1742; apical, 2.35, p=0.0029). We conclude that FF inputs display selectivity for looped neurons in L5, and that this selectivity may result from inputs impinging on the perisomatic dendrites (V1→LM, IT vs. PT), on the apical dendrites (V1→LM, IT vs. IT), or on both dendritic domains (V1→AM, IT vs. IT).

We next measured the connectivity of FB inputs to L5 neurons with different projection targets (Figure 5E). We transfected LM or AM FB axons with ChR2 and recorded different L5 cell types in V1. While perisomatic FB inputs did not distinguish between looped IT neurons and non-looped PT neurons, apical tuft FB inputs were stronger in looped neurons (Figure 5F,G; median ratio of input; looped IT neuron/PT neuron: perisomatic, 1.58; p=0.1465; apical: 3.59, p=0.0134). Consequently, looped IT neurons received a larger fraction of total FB input in their apical compartment compared to PT neurons (Figure 5H), despite having less total dendritic length in L1 (Figure S1). While most active conductances were pharmacologically blocked during our recordings (see Methods), inputs to distal dendritic branches may be filtered and attenuated when measured from the soma due to the passive cable properties of dendritic arbors ^26,27^. However, simulations showed that the weaker sCRACM responses in distal tufts of PT neurons cannot be explained by differences in passive dendritic filtering between the two cell types. In the absence of connectional selectivity, the thicker apical shafts and richer apical tuft arborization of PT neurons (Figure S1) predict that distal FB input arriving at the soma would be larger, not smaller (Figure S2). We also confirmed that the weaker distal FB input in PT neurons was not due to different levels of hyperpolarization-activated current (*I*_*h*_) between the two cell types^4^, since looped IT neurons still received stronger FB input in the apical tuft when measured in the presence of *I*_*h*_ blockers (Figure S3). Thus, the stronger sCRACM signals detected in L1 reflect synaptic selectivity for the terminal tufts of looped IT neurons. We then compared the strength of LM→V1 and AM→V1 FB inputs to looped versus non-looped L5 IT neurons (Figure 5E–H). We again found that FB innervated looped neurons more strongly in the apical compartment, but not in the basal compartment (median ratio of input; looped IT neuron/non-looped IT neuron: perisomatic, 1.29, p=0.5876; apical, 2.24, p=0.0541). As a result, looped neurons received a larger fraction of total FB input on their apical dendrites in comparison to non-looped IT neurons (Figure 5H; looped, 20.8%, non-looped 10.2%, p=0.0009). The stronger apical input to looped neurons could not be explained by differences in dendritic filtering. Firstly, LM- and AM-projecting populations had similar apical dendritic morphologies, with slender tufts that were indistinguishable in L1 (Figure S1). Secondly, the cell type receiving strongest distal input switched from LM-projecting neurons in the LM→V1 experiment to AM-projecting neurons in the AM→V1 experiment (Figure 5H; fraction of total input on apical tuft; AM→V1 input, looped IT neurons, 12.3%; non-looped IT neurons, 4.8%, p=0.025; LM→V1 input, looped IT neurons, 25.3%; non-looped IT neurons, 14.3%, p=0.015). We conclude that the apical dendrites of L5 neurons have privileged access to FB axons when the neurons loop back to the source of those axons. Conversely, FB inputs to the basal dendrites of L5 neurons do not favor looped neurons over non-looped IT or PT neurons.

### CC inputs do not selectively innervate looped L2/3 neurons

Finally, we examined whether CC projections to supragranular neurons would also exhibit a preference for looped connectivity. Unlike FF connections to infragranular neurons, FF input from V1 was equally strong in looped and non-looped L2/3 IT neurons in LM (Figure 6A–C; median ratio of input; looped IT neuron/non-looped IT neuron: 1.12, p=0.6221). Similarly, we measured the strength of AM and LM FB inputs to different L2/3 IT neurons in V1. Total input was indistinguishable between looped and non-looped neurons, either when taking both sources of FB together (Figure 6D–F; median ratio of input; looped IT neuron/non-looped IT neuron: 1.22, p=0.3176) or each individually (median ratio of input; AM→V1 input, looped IT neuron/non-looped IT neuron: 1.22, p=0.3303; LM→V1 input, looped IT neuron/non-looped IT neuron: 1.22, p=0.7173). In summary, in contrast to CC innervation of infragranular neurons, neither FF nor FB inputs preferentially innervate looped neurons over non-looped neurons in L2/3.

**Figure 6.**
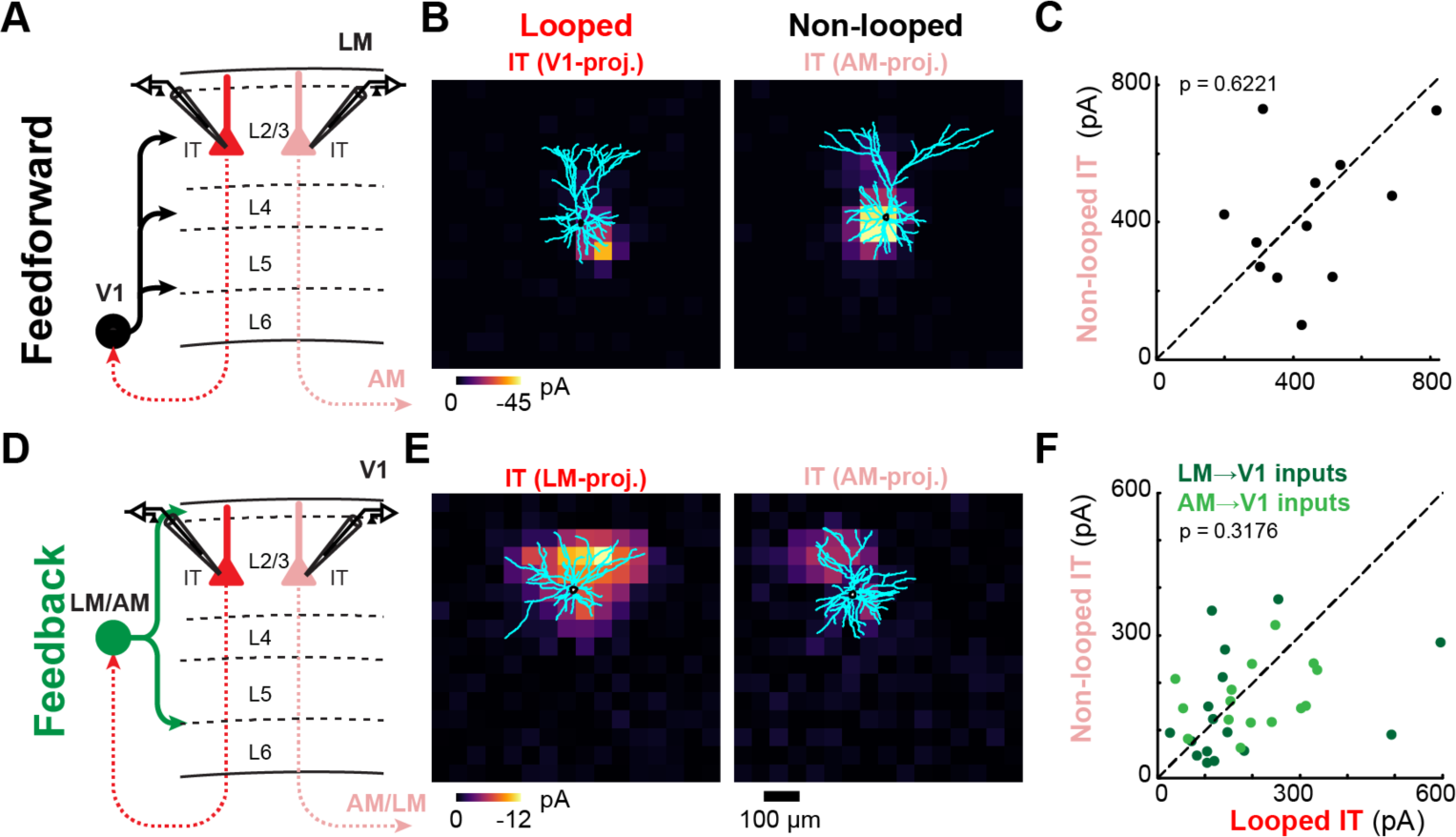
FF and FB connections do not selectively target looped L2/3 neurons. **A**, Configuration of experiment comparing strength of V1 FF input to pairs of looped and non-looped L2/3 IT neurons in LM. **B**, Example pair of sCRACM maps overlaid on reconstructed dendrites. **C**, Paired comparisons of total FF input to looped vs. non-looped IT neurons in L2/3 (n=12 pairs). Statistics, two-tailed Wilcoxon signed-rank test for paired samples. **D**, Configuration of experiment comparing strength of LM or AM FB input to pairs of looped and non-looped L2/3 IT neurons in V1. **E**, Example pair of sCRACM maps overlaid on reconstructed dendrites. **F**, Paired comparisons of total FB input to looped vs. non-looped IT neurons in L2/3 (n=31 pairs; dot colors denote the photostimulated inputs: LM→V1, n=16; AM→V1, n=15).

## Discussion

We comprehensively measured the connectivity of several ascending and descending projections to the three major classes of cortical projection neurons across layers and across areas. We found that both FF and FB preferentially innervated looped over non-looped neurons in the infragranular layers, but not in the supragranular layers. Furthermore, by mapping the dendritic locations of CC synaptic inputs, we show that targeting of looped neurons is often highly subcellular, with FB projections showing selectivity for the apical domains of looped neurons, but not for their basal domains.

### FF and FB connections have similar connectivity rules

In any given layer, FF and FB inputs to projection neuron classes followed similar connectional principles (Figure 7), despite their very different laminar innervation patterns (Figure 1C,D). Given the distinct gene-expression profiles of IT, PT and CT neurons ^22^, the wiring of FF and FB axons could be guided by cell-type specific molecular cues. However, while the cell-type specificity of FF and FB projections was the same, the underlying subcellular specificity differed across L5 cell types (Figure 5 and Figure 7). FB selectivity was always restricted to the apical tufts of L5 neurons, whereas FF selectivity sometimes involved perisomatic dendrites. Thus, FF and FB connectivity could be shaped by a combination of molecular signaling and dendrite-specific plasticity rules ^28,29^, resulting in similar cell-type specificities, but involving different dendritic compartments.

**Figure 7.**
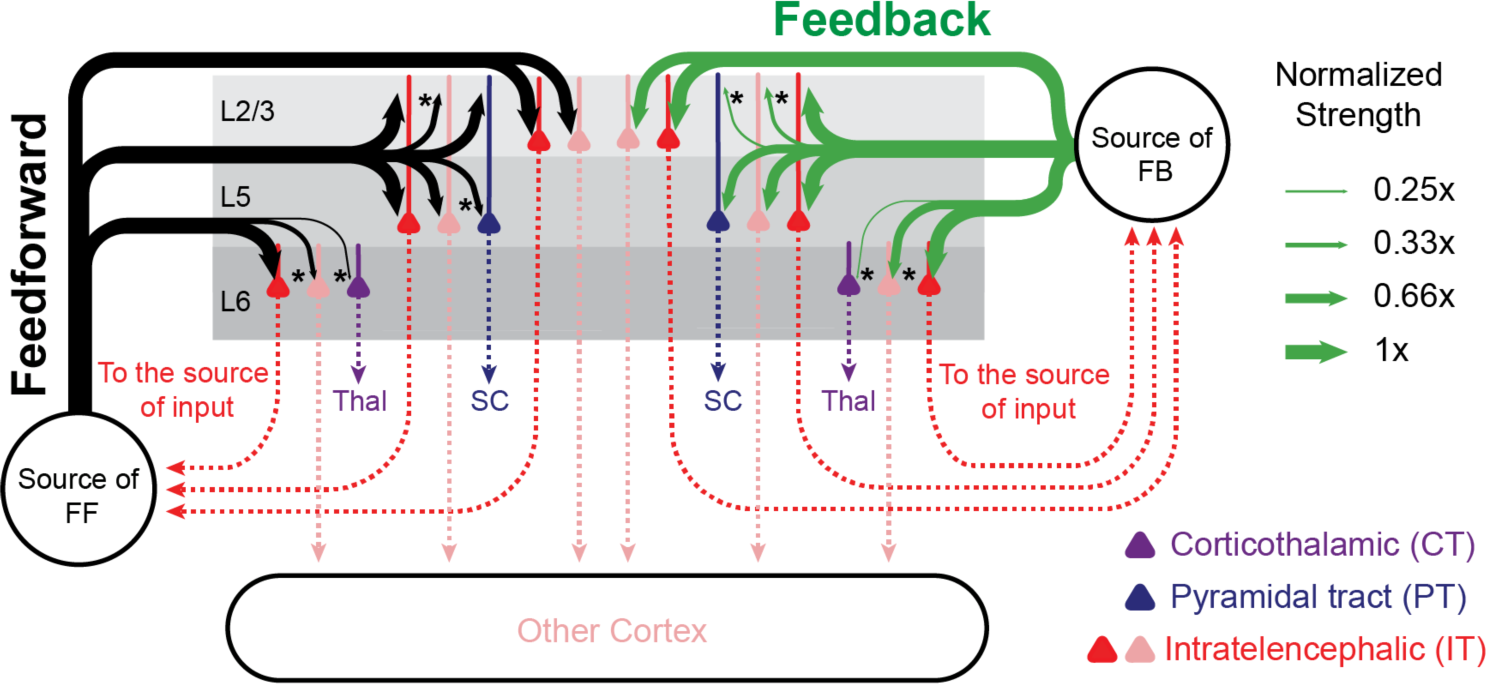
Summary of FF and FB connectivity. The strength of FF and FB inputs to the different cell types is represented by arrow thickness, which is normalized to the looped IT population in each cortical layer. The top and bottom arrows to L5 neurons indicate inputs to apical and perisomatic domains, respectively. Asterisks signify significantly different input strength from the looped population.

### Looped connectivity in CC interactions

Both FF and FB inputs always innervated looped IT neurons more strongly than neighboring subcortical-projecting PT and CT neurons. Input strength was also always higher in non-looped IT neurons versus PT and CT neurons (Figure 7). Thus, the primary cell type targeted by IT-derived CC projections from visual areas are other IT neurons, especially those in deep layers forming a monosynaptic loop with the projection source. These findings accord with rabies virus (RV)-mediated trans-synaptic retrograde tracing experiments reporting that V1 IT neurons receive a larger proportion of their monosynaptic inputs from higher visual cortices than do V1 PT neurons ^30^. They are also consistent with studies showing that monosynaptic FF inputs from primary sensory cortices selectively innervate looped neurons in frontal cortical regions ^31,32^. However, single-cell RV tracings in V1 L6 found that CT neurons had a greater fraction of presynaptic partners in higher-order cortical areas than IT neurons ^33^, and CC fibers are generally considered to be a major synaptic drive of CT neurons ^4,34^. Our functional measurements challenge that view, as we found that CC inputs to CT neurons were relatively weak, with FF and FB connections being ~7- and ~10-fold weaker, respectively, than those in looped IT neurons (Figure 4 and Figure 7). Thus, IT neurons are the principal excitatory targets of CC visual afferents in all layers, and looped IT neurons are the main recipients of these afferents in deep layers. The predominance of direct, monosynaptic excitatory loops in interareal connections could be an organizing principle of long-range CC communication, performing a conserved computational motif across functionally divergent cortical areas.

The specificity of CC connections for looped neurons in infragranular layers was not absolute, as we detected monosynaptic CC inputs in all cell types, consistent with previous reports ^30,35^. However, our measurements likely underestimate the selectivity of CC inputs for looped neurons. Firstly, many, if not most, projection neurons target multiple cortical areas ^21,36^. Thus, since retrograde tracer injections only label a fraction of neurons projecting to the injection site, some non-looped neurons might in fact send a looped projection despite not being retrogradely labeled, thereby reducing our observed specificity for looped neurons. Secondly, looped connectivity would also be underestimated if it only involves CC afferents originating from a specific layer, as we expressed ChR2 non-specifically in projection neurons spanning multiple layers. Future experiments with layer-specific transgenic mouse lines ^19^ will make it possible to compare the specificity of projections with different laminar origins.

FF and FB connections to L2/3 neurons were unusual in that they did not show a predilection for forming monosynaptic loops (Figure 6, Figure 7). While the axons of most L2/3 neurons branch to innervate multiple cortical areas ^21^, the absence of loop specificity in L2/3 is unlikely to be caused by looped neurons being misattributed as non-looped neurons due to incomplete labeling from tracer injections. This is because L5 showed looped connectivity despite having a larger percentage of neurons with bifurcating axons (Figure 1).

It remains unknown whether the CC loops that we unveil here arise from pairs of individual neurons in different cortical areas directly targeting each other in a recurrent loop. Given that ChR2 was expressed in multiple presynaptic neurons, our experiments measured the selectivity of afferent populations and do not have the resolution to resolve interareal loops with single-cell resolution. There is evidence that L5/6 cortical neurons receive long-range CC inputs preferentially from L5/6 neurons ^35^. Moreover, L6 IT neurons innervate deep layers in their target areas ^22^, and L5 IT neurons have access to corresponding L5 IT neurons in distant cortical regions, as indicated by their axonal arborization pattern ^19,22^. Thus, while our findings are consistent with pairs of deep-layer neurons in different cortical areas engaging in bidirectional monosynaptic loops, the prevalence of such a circuit arrangement has yet to be determined.

### A role for the apical tufts of L5 neurons in looped hierarchical interactions

FB strength in distal tufts of L5 neurons was relatively larger in looped neurons versus non-looped PT or IT neurons (Figure 5). Given also that FB strength in basal dendrites of looped and non-looped L5 neurons was indistinguishable, these results point to a central role for the apical dendritic trees of L5 IT neurons in looped hierarchical interactions. FB inputs to apical tufts might be selectively involved in recurrent computations, while perisomatic inputs might mediate other hierarchical exchanges. We found that, on average, the apical dendrites of looped L5 neurons were also preferentially targeted by FF axons, even when their basal dendrites were not. However, subcellular selectivity was inconsistent across the two FF projections studied (Figure 5), suggesting that for FF pathways apical dendritic targeting plays a less prominent role in looped interactions. Through a dynamic interplay of active and passive membrane properties, the apical arbors of L5 pyramidal neurons can perform complex computations combining inputs from different dendritic compartments ^26,37,38^. The apical dendrites of L5 IT neurons are less well understood than those of PT neurons, being thinner and less experimentally tractable. Understanding hierarchical computations will require elucidating how looped L5 IT neurons integrate CC inputs in their distal tufts with inputs arriving in other dendritic regions.

### Functional implications

The preferential innervation of looped neurons by both FF and FB afferents supports several theories advocating looped computations in CC circuitry ^5–7,9–11,39,40^. Our observations suggest that L5/6 IT neurons might be critical to the implementation of these long-range loops. CC inputs and postsynaptic neurons in their target area have similar tuning properties ^41–43^ and overlapping receptive fields ^44^. Thus, given the prominence of looped connectivity in FF and FB visual pathways, related visual signals are likely relayed back and forth between interconnected neurons located at different hierarchical levels.

The lack of specificity for looped neurons in L2/3 is surprising in light of experimental evidence associating this layer with prediction errors. Neural responses consistent with predictive processes have been identified in L2/3 in several cortical areas ^45–48^ and the layer is considered critical for generating prediction error signals ^5,6,49^. Predictive models require excitatory and inhibitory looped interactions of FB inputs with lower-level neurons to signal violations of expectations ^6^. As our measurements can only detect direct monosynaptic excitatory inputs, FB afferents could still selectively target looped L2/3 neurons polysynaptically via local inhibitory ^45^ or excitatory neurons. However, current circuit-level models of predictive coding may need to consider the fact that populations of CC visual afferents show no predilection for forming monosynaptic excitatory loops in L2/3.

Our finding that the apical dendrites of looped L5 IT neurons are selectively innervated by FB inputs is consistent with distal tufts having a specialized role in processing reciprocal signals. According to recent models, such FB innervation could allow neurons embedded in a hierarchical network to optimize synaptic weights towards the global desired output, akin to the backpropagation algorithm used to train artificial neural networks ^8–11,40^. The apical dendrites of looped neurons are integral to many such models, wherein apical inputs trigger synaptic plasticity of basal inputs to instruct learning. Thus, selective targeting of the apical compartments of looped neurons by descending inputs may allow L5 IT neurons to update synaptic strengths based on activity in the higher-order areas that they project to. Such a role in looped interactions does not negate the involvement of apical dendrites in other non-looped computations. Since L5 PT neurons receive more inputs from frontal and associative areas than IT neurons ^30^, their thicker apical dendrites might be specialized in mediating top-down processes that do not require excitatory looped connectivity, such as brain-state-dependent and attentional modulations of perceptual saliency ^50–52^. Our observations identify L6 and the apical dendrites of L5 as key players in recurrent interareal cortical interactions and provide a framework for future studies on the role of the major classes of projection neurons in different hierarchical processes.

## Materials and Methods

### Animal surgeries

All procedures were reviewed by the Champalimaud Centre for the Unknown Ethics Committee and performed in accordance with the Portuguese Veterinary General Direction guidelines. Surgeries were conducted in either male or female C57BL/6J mice (P26−P28) under anesthesia (intraperitoneal, 37.5 mg/kg ketamine, 0.5 mg/kg medetomidine). Virus expressing ChR2 (AAV-2/1-CAG-Channelrhodopsin-2-Venus, Penn Vector Core, University of Pennsylvania; 20–25 nl) was delivered intracortically either to V1 to label FF projections or LM/AM to label FB projections, and co-injected with red-fluorescent microspheres (Red Retrobeads IX, Lumafluor; 10– 12.5 nl) to retrogradely label cells projecting to the source of FF/FB input. A second retrograde tracer (Cholera toxin subunit B, Alexa Fluor 647 Conjugate, Invitrogen, 50–60 nl, 1.0 mg/mL) was injected elsewhere to label cells projecting to a different cortical or subcortical area. For axonal quantification, AAV-GFP (Penn Vector Core) was used. Pulled glass injection pipettes (Drummond Scientific) had tip diameter of 15–20 μm. Stereotaxic coordinates for the primary visual cortex (V1) and lateromedial area (LM) were measured from the midline and from the posterior-most point of the transverse sinus (lateral of midline/anterior of transverse sinus/depth in mm): V1 (2.3/1.3/0.775), LM (3.5/1.7/0.9). Stereotaxic coordinates for the anteromedial area (AM) and superior colliculus (SC) were measured from the midline and from the sinus confluence, the point at which the transverse sinuses meet the superior sagittal sinus (lateral of midline/anterior of sinus confluence/depth in mm): AM (1.6/1.25/0.8), SC (0.5/0.4/1.5 & 1.8). The accuracy of LM and AM coordinates was verified by intrinsic signal imaging in a subset of animals. Stereotaxic coordinates for the dorsal lateral geniculate nucleus (dLGN) and the lateral posterior nucleus (LP) of the thalamus were measured from the midline and from bregma (lateral of midline/posterior of bregma/depth in mm): dLGN (2.3/1.75/2.8), LP (1.35/1.75/2.65 & 2.85). Animals were maintained at 37°C on a heating pad during surgery and returned to their home cages after surgery (maximum of 5 animals per cage). All animals were housed in a room with a regular 12 h light/dark cycle.

### Slice preparation

14–20 days (age range: P40–P48) after the surgery, mice were decapitated under deep anesthesia (isoflurane) and brains were dissected in ice-cold choline chloride solution (110 mM choline chloride, 25 mM NaHCO_3_, 25 mM D-glucose, 11.6 mM sodium ascorbate, 7 mM MgCl_2_, 3.1 mM sodium pyruvate, 2.5 mM KCl, 1.25 mM NaH_2_PO_4_ and 0.5 mM CaCl_2_ (Sigma); aerated with 95% O_2_/5% CO_2_) and sectioned in 300-μm-thick coronal slices using a Leica VT1200S vibratome. Slices were then incubated for 30 min at 37°C in artificial cerebrospinal fluid (127 mM NaCl, 25 mM NaHCO_3_, 25 mM D-glucose, 2.5 mM KCl, 1 mM MgCl_2_, 2 mM CaCl_2_ and 1.25 mM NaH_2_PO_4_ (Sigma); aerated with 95% O_2_/5% CO_2_).

### Electrophysiology and photostimulation

Neurons were patched with borosilicate pipettes (resistance 3−5 MΩ, Werner Instruments) filled with potassium gluconate intracellular solution (128 mM potassium gluconate, 4 mM MgCl_2_, 10 mM HEPES, 1 mM EGTA, 4 mM Na_2_ATP, 0.4 mM Na_2_GTP, 10 mM sodium phosphocreatine, 3 mM sodium L-ascorbate, 3 mg/mL biocytin (Sigma) and 5 μg/mL Alexa Fluor 488 dye (Thermo Fisher Scientific); pH 7.25, 290 mOsm). All recordings were performed at room temperature (22−24°C) and with the presence of TTX (1 μM), 3-((R)-2-carboxypiperazin-4-yl)-propyl-1-phosphonic acid (CPP, 5 μM) and 4-aminopyridine (4-AP,100 μM) in the bath. For Figure S3, ZD7288 (10 μM) was also applied to control for *I*_*h*_ differences between cell types. Areas LM and AM were identified by the presence of Venus-expressing FF axons. For sCRACM mapping, fluorescent-positive cells were recorded sequentially in voltage clamp (−70 mV) at depths of >30 μm in the same slice. Double-labeled cells were not recorded. A blue laser (473 nm, Cobolt Laser) was used for photostimulation to evoke excitatory postsynaptic currents (EPSCs). Duration (1 or 4 ms) and intensity (0.1−1.1 μW) of light pulses were controlled with a Pockels cell (ConOptics) and a shutter (Thorlabs). The laser beam was rapidly repositioned using galvanometer mirrors (Thorlabs) and delivered through an air immersion objective on either a 16 x 16 grid (L2/3 cells) or a 12 x 24 grid (L5 and L6 cells) with 50-μm spacing and 400 ms inter-stimulus interval. Stimuli were given in a spatial sequence pattern designed to maximize the time between neighboring locations. The stimulus pattern was flipped and rotated between maps to avoid sequence-specific responses. sCRACM maps were repeated 2–5 times for each cell. Laser power was manually adjusted in each experiment using a graduated neutral density filter (Edmund Optics) so that peak amplitudes smaller than 100 pA were evoked in the most excitable locations for the first recorded cell in the pair. Pairs of different projection neurons recorded at similar cortical depths in the same layer and in close proximity (mean ± s.d.: 73.27 ± 47.41 μm) were photostimulated using the same laser power and pulse duration. The order in which cell types were recorded alternated between pairs. Data were acquired with a Multi clamp 700B amplifier (Axon Instruments), and digitalized with National Instruments acquisition boards controlled under Matlab using Ephus ^53^.

### Immunohistochemistry and dendritic reconstructions

After whole-cell recordings, biocytin-filled neurons were fixed overnight in 4% paraformaldehyde (PFA) at 4°C and transferred to phosphate-buffered saline the following day. Prior to staining, slices were rinsed in phosphate buffer (PB) 0.1 M. Endogenous peroxidases were quenched with 1% H_2_O_2_ (Sigma) in PB 0.1 M for 45 min at room temperature. Slices were rinsed again in PB 0.1 M and incubated in the ABC reaction (Vector Laboratories) for ~12 h at room temperature (22−24°C). After successive PB 0.1 M and Tris-buffered saline (TBS) washings, slices were subjected to the diaminobenzidine (DAB) reaction for 30−50 min (using 30 mL of TBS, 90 μL 3% H_2_O_2_, 225 μL of NiCl_2_ (250 mM) and 7 mg of DAB (Sigma)). The DAB reaction was stopped with TBS. Slices were mounted and coverslipped with Mowiol mounting medium. Dendrites were reconstructed with Neurolucida software (MBF Bioscience) using the x40 magnification objective lens of an Olympus BX61 microscope. Tracings were imported into Matlab, corrected for shrinkage and analyzed using custom routines. Dendritic length density was calculated in 50-μm bins and interpolated for display.

### Laminar distribution of projection neuron subtypes and FF/FB axons

Injected animals were intracardially perfused with 4% PFA 14 days post-surgery and cryostat-sectioned in 20-μm thick coronal slices. Slices were stained for DAPI and imaged with the x20 objective of a Zeiss AxioImager M2 widefield fluorescence microscope. For quantification of different projection neurons in V1/LM cortex (Figure 1A,B), 3 animals were used per dataset (8 slices per animal). Cells were counted within a 1000 x 1000 μm (V1) or 600 x 1000 μm (LM) area. To normalize cell depth, fractional cortical depth (pia–cell distance/pia– white matter distance) was multiplied by average cortical thickness across the 8 slices. For quantification of axons (Figure 1C,D), 3 animals were analyzed for both FF and FB datasets (8 slices per animal). Vertical fluorescence profiles of GFP-expressing axons were measured using ImageJ after subtracting background fluorescence.

### Data analysis

The boundaries between layers were established as L1: pia−100 μm; L2/3: 100−350 μm; L4: 350−450 μm; L5: 450−650 μm; L6 650−950 μm. To correct for differences in cortical thickness due to variability in slicing angle of brain sections, the fractional cortical depth of each recorded neuron was positioned on a reference cortical slice with a thickness of 950 microns. Only pairs of cells with an intersomatic distance of <200 μm were included in analyses. In cases where multiple cells were recorded in the same slice, cells nearest each other were paired. Pairs in which neither cell showed detectable input (total sCRACM input < 5 pA) were discarded. For L5 datasets, pairs with any cut apical dendrites were also discarded. For inclusion in fraction-to-apical analyses, both cells in the pair required detectable input. Statistical comparisons were made using the two-tailed Wilcoxon signed-rank test for paired samples. All statistical analyses were performed with Matlab.

### Simulations of passive dendritic filtering of L1 inputs

Traced dendrites of L5 neurons in V1 were imported into the NEURON simulation environment ^54^. We set the diameter of the somata, apical trunks and apical tuft branches for the different projection neurons using manual measurements from biocytin-stained arbors (Figure S1). For each cell belonging to a given projection type, segments from the same dendritic compartment were assigned the same diameter value (apical trunk width, μm: SC-projecting, 1.7, AM-projecting, 1.14; LM-projecting, 1; apical tuft branch width, μm: SC-projecting, 0.7, AM-projecting, 0.4, LM-projecting, 0.43). The biophysical properties used for simulations were as follows: cytoplasmic resistivity, R_a_=35.4 cm; specific membrane capacitance, C_m_=1 F/cm^2^; resting conductance, g_pas_=1/20000; resting potential, E_pas_=-65 mV. We applied a synaptic density of 0.2 synapses per μm of apical dendritic segment when distributing passive synapses over the apical tree. The location of synapses along individual dendritic segments was randomly determined, and synaptic conductance was approximated by an alpha function, with parameters τ = 0.1 ms and g_max_=1 μS. For each neuron, we then simulated responses to apical tuft inputs under single-electrode voltage-clamp at the soma (−70 mV, 10 MΩ resistance), assuming no axonal selectivity for postsynaptic cell types. We conducted 100 simulations for each neuron and quantified the mean charge measured at the soma (Figure S2).

## Acknowledgments

We thank N. Yamawaki, B. Atallah and M. Fridman for critical comments on the manuscript. This work was supported by a fellowship from Fundação para a Ciência e a Tecnologia to HY and by grants from Marie Curie (PCIG12-GA-2012-334353), la Caixa Banking Foundation (LCF/PR/HR17/52150005) and Fundação para a Ciência e a Tecnologia (LISBOA-01-0145-FEDER 030328 and Congento LISBOA-01-0145-FEDER-022170), co-financed by FCT (Portugal) and Lisboa2020, under the PORTUGAL2020 agreement (European Regional Development Fund), and the Champalimaud Foundation.

