## Supplementary Figures for "Laminar-specific cortico-cortical loops in mouse visual cortex"

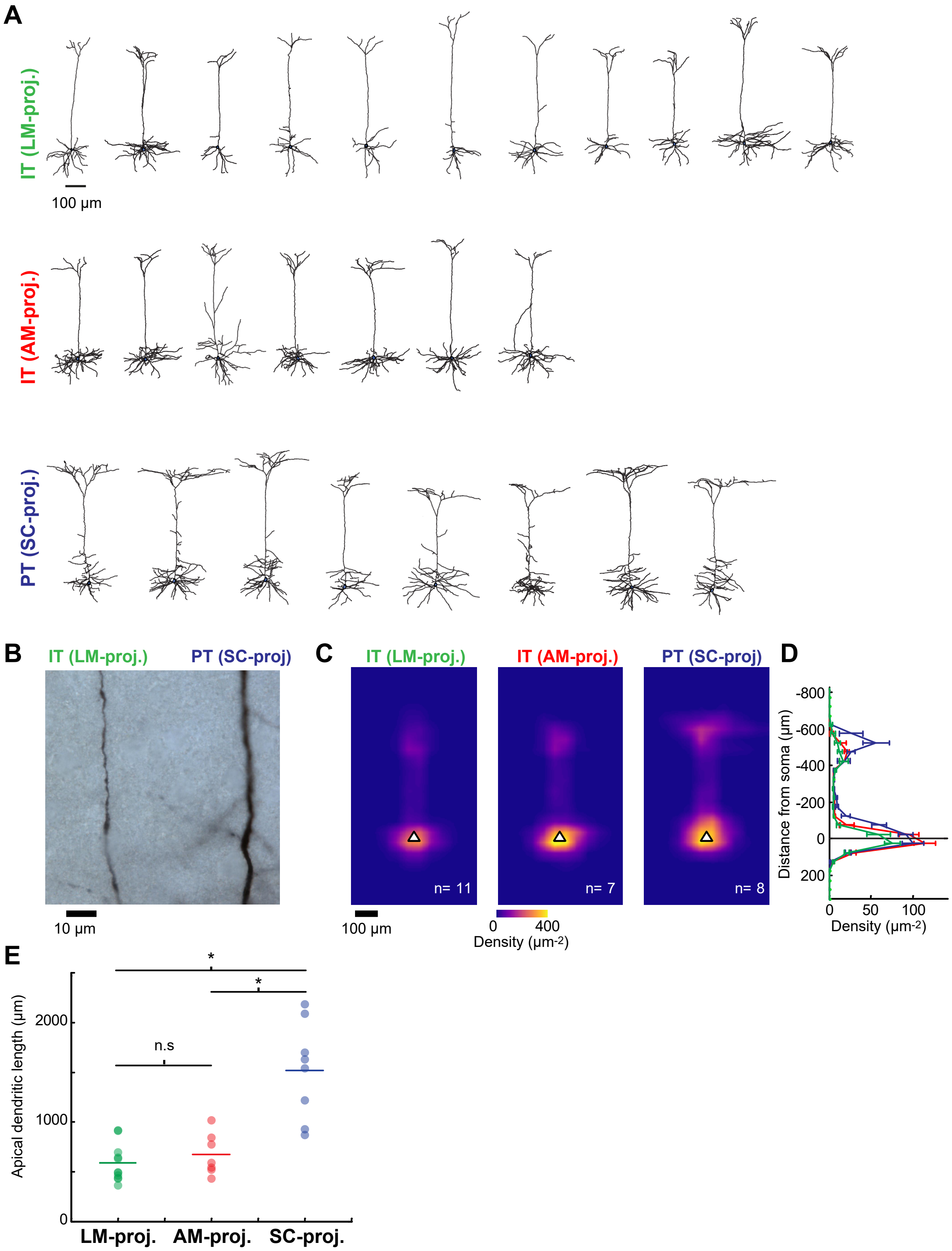


**Figure S1. Dendritic morphology of the different L5 projection neuron types in V1.**

(A) Reconstructed dendritic morphologies of the three different L5 projection neurons recorded in V1. Top and middle, IT neurons projecting to LM or AM; Bottom, PT neurons projecting to SC.

(B) Brightfield image showing representative example of apical shaft segments from a pair of biocytin-stained IT and PT L5 neurons. The apical dendrites of SC-projecting PT neurons were of larger diameter than those of same-layer LM- and AM-projecting IT neurons.

(C) Average normalized dendritic length density of the three cell types, aligned by soma position (white triangle).

(D) Mean vertical profiles of dendritic length density (error bars, s.e.m).

(E) Total apical tuft dendritic length for the three cell types. Apical tuft branches of SC-projecting PT neurons are more extensive than those of LM- and AM-projecting IT neurons (Kruskal-Wallis test followed by Tukey-Kramer HSD post-hoc test, SC-projecting vs. LM-projecting, p=0.0006; SC-projecting vs. AM-projecting, p=0.0168; LM-projecting vs. AM-projecting, p=0.8044).

.

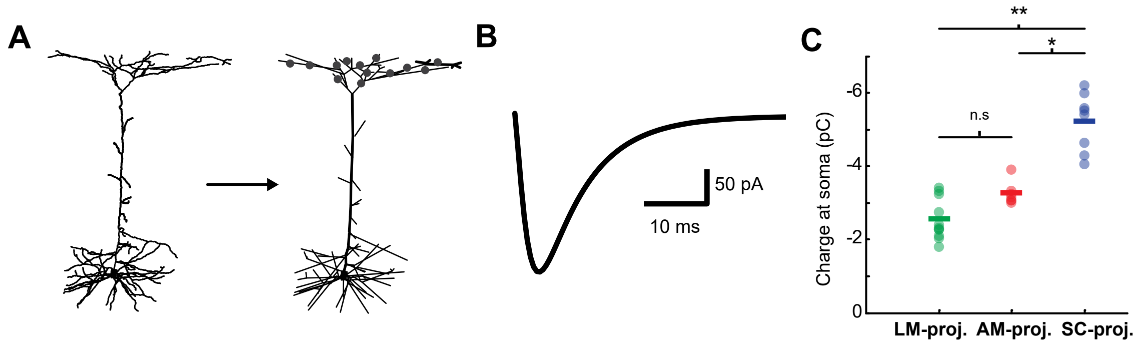


**Figure S2. Simulations of the dendritic filtering of distal apical inputs**.
(A) Example dendritic simulation of a L5 neuron. Reconstructed dendritic arbors were imported into the NEURON environment. Synapses were randomly placed with constant density along apical tuft dendritic segments.
(B) Simulated EPSC at the soma evoked by L1 input under voltage-clamp conditions for the example neuron shown in A.
(C) Mean somatic charge per cell (based on 100 simulations) resulting from apical tuft input across the three projection neuron populations. Apical inputs lacking cell-type selectivity and exhibiting equal synaptic density across the different cell types generate larger somatic currents in SC-projecting neurons versus AM- or LM-projecting neurons (Kruskal-Wallis test followed by Tukey-Kramer HSD post-hoc test, SC-projecting vs. LM-projecting, p=0.0001; SC-projecting vs. AM-projecting, p=0.0325; LM-projecting vs. AM-projecting, p=0.3626).

**Young et al.**
**SFigure 2**

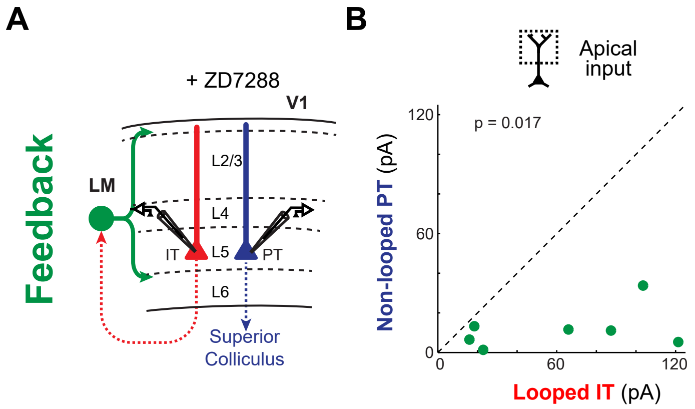


**Figure S3. FB input to looped L5 IT neurons versus PT neurons in the presence of *Ih* blockers.**
(A) Configuration of experiment comparing strength of LM FB input to pairs of looped IT and non-looped PT neurons in V1 L5. ZD7288 was added to the bath solution to block *Ih* currents.
(B) Paired comparisons of apical dendritic FB input to looped IT neurons vs. non-looped PT neurons in L5 (n=7 pairs).
